# Conditioning electrical stimulation fails to enhance sympathetic axon regeneration

**DOI:** 10.1101/2023.02.03.527071

**Authors:** Tina Tian, Kevin Patel, David Kim, HaoMin SiMa, Alandrea R. Harris, Jordan N. Owyoung, Patricia J. Ward

## Abstract

Peripheral nerve injuries are common, and there is a critical need for the development of novel treatments to complement surgical repair. Conditioning electrical stimulation (CES) is a novel variation of the well-studied perioperative electrical stimulation treatment paradigm. CES is a clinically attractive alternative because of its ability to be performed at the bedside prior to a scheduled nerve repair surgery. Although 60 minutes of CES has been shown to enhance motor and sensory axon regeneration, the effects of CES on sympathetic regeneration are unknown. We investigated how two clinically relevant CES paradigms (10 minutes and 60 minutes) impact sympathetic axon regeneration and distal target reinnervation. Our results indicate that the growth of sympathetic axons is inhibited by CES at acute time points, and at a longer survival time point post-injury, there is no difference between sham CES and the CES groups.

We conclude sympathetic axons may retain some regenerative ability, but no enhancement is exhibited after CES, which may be accounted for by the inability of the electrical stimulation paradigm to recruit the small-caliber sympathetic axons into activity. Furthermore, 10-minute CES did not enhance motor and sensory regeneration with a direct repair, and neither 60-minute nor 10-minute CES enhanced motor and sensory regeneration through a graft. Further studies will be needed to optimize electrical stimulation parameters to enhance the regeneration of all neuron types.

## INTRODUCTION

Peripheral nerve injuries (PNIs) are common and can lead to lifelong functional deficits.^1–4^ The current standard of care relies on the inefficient process of spontaneous axon regeneration after surgical decompression or surgical repair with only 10% of PNI patients ever achieving full functional recovery.^5^ Therefore, there is a critical need for the development of novel interventions that complement surgical repair.

One hour of electrical stimulation (ES) of the proximal stump at the time of nerve repair or decompression is a well-studied experimental intervention that enhances motor and sensory axon regeneration.^6–12^ Previous research on ES applied the intervention perioperatively which improved outcomes in animal models and in clinical trials of carpal tunnel syndrome, cubital tunnel syndrome, shoulder function after oncologic neck dissection, and digital nerve transection.^7,13–17^

The application of 60 minutes of ES *prior* to injury for nerve conditioning, known as conditioning ES (CES), is a novel way that may also improve outcomes after PNI.^18–25^ For nerve injury cases in which surgical interventions are delayed, such as those involving nerve transfers, CES in the clinic prior to a scheduled operation may be beneficial.^23,24^ One hour of CES 7 days prior to nerve injury and repair improved motor and sensory axon regeneration in rat models.^23,24^ CES was also more favorable than perioperative ES (PES) or a combination of CES and PES in a rat model of tibial nerve autograft repair.^19^ More recently, a 10-minute PES session has been described as being similarly effective as a 60-minute PES session at enhancing axon regeneration,^26,27^ but 10-minute CES has not been tested head-to-head with 60-minute CES. To date, CES has primarily been investigated by a single research group, and its promise should be replicated by independent groups and with varying repair strategies.

Motor axons have understandably been the major focus of regeneration-related studies, yet peripheral nerves are primarily composed of sensory and sympathetic axons. Only 6% of the rat sciatic nerve is composed of myelinated motor axons while unmyelinated sympathetic axons make up 23%.^28^ Notably, neither a conditioning lesion nor ES has been shown to promote sympathetic axon elongation,^29–32^ and the effect of CES on sympathetic axon regeneration is currently unknown. Herein, we compared the outcomes of two clinically relevant CES paradigms (10 minutes and 60 minutes) on sympathetic axon regeneration and long-term sympathetic muscle reinnervation. The regeneration of sympathetic axons to muscle should be a priority because sympathetic signaling at neuromuscular junctions (NMJs) provides trophic support and stability to the synapse as well as maintains muscle mass.^33–37^ These findings highlight the importance of functional sympathetic axon regeneration in the context of nerve injury, and importantly, muscle atrophy is a clinically relevant characteristic of reinnervated muscle.^35–37^ Because of the abundance of evidence that ES enhances motor and sensory regeneration, we hypothesized that CES will enhance sympathetic regeneration. However, our results did not support this hypothesis and also revealed important findings regarding motor and sensory regeneration with CES.

## METHODS

### Animals

Experiments were conducted on adult (2–5 months old) mice weighing 18–30 g using approximately equal males and females for each assay. Mice were randomly assigned to sham CES, 10-minute CES, or 60-minute CES groups. Tg(Thy1-YFP)16Jrs transgenic mice (Jackson laboratory stock no. 003782, “YFP-16”) or wild type mice C57BL/6J (Jackson laboratory stock no. 000664) were used in these experiments. The YFP-16 mice allowed us to differentially visualize regenerating motor and sensory axons versus sympathetic axons as the yellow fluorescent protein is robustly expressed in somatic motor neurons and a small subset of primary sensory neurons but has negligible expression in postganglionic sympathetic neurons,^38,39^ which can henceforth be visualized with immunohistochemistry. Experimental groups with final numbers of mice are presented in **Table 1**.

**Table 1:** Detailed breakdown of experimental groups. The n represents the final number of animals included in each analysis. *1-2 animals were excluded due to poor quality nerve sections.

| Experiment | Treatment | n = | Strain |
| --- | --- | --- | --- |
| Axon regeneration through a nerve graft (1 week) | Sham CES | 2 males<br>3 females* | Tg(Thy1-YFP)16Jrs |
|  | 10-minute CES | 2 males<br>3 females* | Tg(Thy1-YFP)16Jrs |
|  | 1-hour CES | 1 male<br>4 females* | Tg(Thy1-YFP)16Jrs |
| Axon regeneration with direct nerve repair (2 weeks) | Sham CES | 4 males<br>3 females* | Tg(Thy1-YFP)16Jrs |
|  | 10-minute CES | 3 males*<br>4 females | Tg(Thy1-YFP)16Jrs |
|  | 1-hour CES | 2 male<br>4 females* | Tg(Thy1-YFP)16Jrs |
| Long-term target reinnervation and retrograde labeling (10 weeks) | Sham CES | 4 males<br>4 females | C57BL/6J |
|  | 10-minute CES | 5 males<br>4 females | C57BL/6J |
|  | 1-hour CES | 4 males<br>4 females | C57BL/6J |

Some animals were excluded from analysis in the 1- and 2-week regeneration assays due to poor sciatic nerve section quality. For a nerve section to be included in analysis, the imaged section must include the injury site, and the distal stump must extend at least 2500 µm from the injury site. If one or more of these criteria were not met, then the section was excluded. If a sciatic nerve did not have at least 2 sections that met the criteria for inclusion, the animal was excluded from analysis. All experiments were approved by the Institutional Animal Care and Use Committee of Emory University (PROTO202100177). A summary of methods is presented in **Figure 1**.

**Figure 1:**
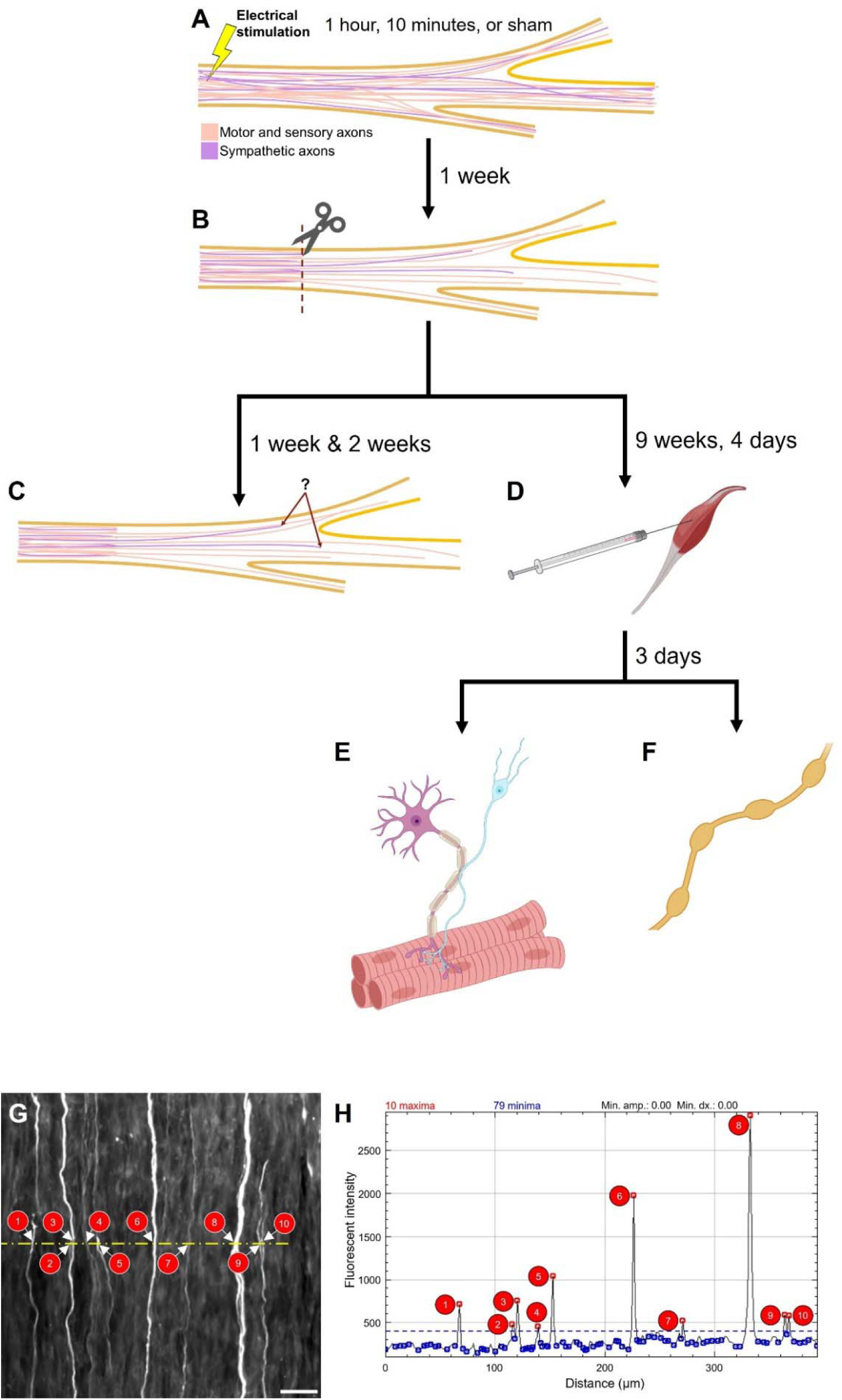
Summary of methods. (A) Sciatic nerves were sham stimulated or electrically stimulated for 1 hour or 10 minutes. (B) One week later, the nerves were transected and repaired with a sciatic nerve graft from a littermate or simply transected and repaired with fibrin glue. (C) Axon profiles were counted 1 week after nerve graft repair and 2 weeks after direct repair. (D) In the long-term reinnervation group, their gastrocnemii muscles were injected with cholera toxin subunit B Alexa Fluor 555 at 9 weeks and 4 days after direct nerve repair. (E) Three days post-injection, the mice were perfused. Their gastrocnemii muscles were harvested for evaluation of the presence of catecholamines at the neuromuscular junction, (F) and their L2-L5 lumbar sympathetic ganglia were harvested for retrogradely-labeled neuron counts. (G) Manually counted axon profiles at a linear region of interest (ROI, dotted yellow line) across a nerve section correlated with (H) maxima on the ROI’s plot profile with maxima threshold set at 400 (dotted blue line). Screenshot of the plot profile window displays the number of maxima in the top left. Scale bar = 50 µm. *D-F created with BioRender.com*.

### Surgical procedures

Anesthesia was induced with 3% isoflurane in 1 L/min oxygen and maintained at 2% isoflurane in 1 L/min oxygen. The sciatic nerve was exposed via a small (∼1 cm) skin incision along the femur followed by blunt dissection between the hamstring muscles using straight scissors (Fine Science Tools, 14084-09). Two monopolar needle electrodes (Ambu, #74325-36/40) were placed on either side of the sciatic nerve and were used to deliver electrical stimulus pulses to the nerve. Using our custom Labview® software,^40^ which allows for both cost efficiency and customizability of various laboratory stimulation paradigms, electrical stimulation (20 Hz for one hour or 10 minutes, 0.1 ms duration pulses) was applied at a supramaximal intensity.^11,41–43^ Supramaximal intensity was calculated as 1.5-times the voltage required to evoke a maximal muscle response (typically 3-4 V). Sham animals had their sciatic nerves exposed with electrodes placed for 60 minutes but were not stimulated. In the 10-minute CES group, after 10 minutes of ES, the stimulation was turned off while the electrodes remained in place for another 50 minutes. All mice were anesthetized for equal amounts of time. The muscle was closed with 5-0 absorbable suture (CP Medical, 421A), and the skin was closed with 5-0 non-absorbable suture (MYCO Medical Supplies, N661-SI) **(Figure 1A)**. Seven days later, mice were re-anesthetized, and the sciatic nerve was transected with sharp microscissors (Fine Science Tools, item no. 15006-09). It was then either repaired with a nerve graft at least 5 mm long from a wild type cagemate for 1 week acute axon regeneration studies (which allows for a dark background for YFP+ axons to regenerate into) or repaired by simple end-to-end anastomosis using fibrin glue, composed of fibrinogen and thrombin in a 2:1 ratio, for 2-week acute axon regeneration studies and long-term reinnervation studies **(Figure 1B)** as we have previously described in detail.^41,44,45^ Animals received 5 mg/kg of 1.5 mg/ml oral suspension meloxidyl (Pivetal, ANADA 200-550) prior to surgery and then once a day for 3 days postoperatively for pain control.

Mice were euthanized at 1, 2, or 10 weeks post-PNI with an intraperitoneal (IP) injection of 150 mg/kg 10 mg/ml Euthasol® solution (0.75ml Euthasol® [Virbac AH Inc, Fort Worth, TX, ANADA 200-071, NDC 051311-050-01] in 29.25 ml 0.9% bacteriostatic sodium chloride [Hospira, Inc., Lake Forest, IL, NDC 0409-1966-02]). Upon verification of no response to noxious stimuli, mice in the 10-week post-PNI group were exsanguinated with 0.9% NaCl (Fisher Scientific, S2711) and then perfused with 4% paraformaldehyde (Sigma-Aldrich, P6148) in 0.01M phosphate buffered saline (1X PBS, Sigma-Aldrich, P4417.

Notes on the choice of experimental models: Nerve grafts are frequently used in clinical scenarios in which CES would be useful. Thus, in one assay we measured axon elongation into grafts after 1 week of growth. A longer survival time is necessary to measure the reinnervation of peripheral targets, so we opted to perform CES followed by transection and immediate repair. If we had used a graft, the regenerating axons would need to cross 2 coaptation sites instead of one, which could make results more difficult to interpret.

### Axon profile quantification

Sciatic nerves were collected 1 week post-nerve grafting or 2 weeks post-nerve direct repair while mice were under anesthesia and postfixed for 30 minutes with 4% paraformaldehyde in 1X PBS **(Figure 1C)**. Nerves were then placed in 20% sucrose with 0.02% sodium azide (Fisher Scientific, S227I-25) for cryoprotection and allowed to sink at 4°C prior to being cryosectioned at 20 µm and placed on charged slides (VWR Superfrost® Plus, 48311-703).

Sciatic nerve sections were blocked with 10% normal goat serum (VWR International, GTX73206) in 1X Tris-buffered saline (Pierce, 28358), 0.1% Tween® 20 detergent (TBST) (Fisher Scientific, BP337500) for 1 hour at room temperature (RT). Rabbit anti-tyrosine hydroxylase (TH, 1:750, Abcam Ab112) was used to visualize sympathetic axons as TH is the rate-limiting enzyme for norepinephrine, the major neurotransmitter of postganglionic sympathetic neurons, synthesis.^46^ This primary antibody was diluted in the blocking buffer and allowed to incubate at RT in a humidity chamber overnight followed by 3 10-minute washes with TBST. Secondary antibody goat anti-rabbit Alexa Fluor™ 647 (1:200, Invitrogen A21245) in blocking buffer was allowed to incubate at RT for 2 hours followed by 3 10-minute washes with TBST. Dried slides were then mounted with Fluoro-Gel with Tris (Electron Microscopy Sciences, 17985-10) and imaged on a Nikon A1R confocal microscope on the 20x objective on the resonant scan mode, 8x averaging. The entirety of the nerve section was imaged with the “ND Acquisition + Large Images” wizard.

In Fiji, linear regions of interest (ROIs) were created that spanned the width of the nerve sections every 500 µm distally, starting from the injury site using a custom-made macro. One linear ROI was created 250 µm proximal to the injury site and used as the control for the calculation of regenerative indices. The mean fluorescent intensities of lightly fluorescent YFP+ and TH+ axons were measured to identify a threshold for each nerve section. At each ROI, a “Plot Profile” was generated, after which the “Find Peaks” function under Broadly Applicable Routine (BAR) was utilized.^47^ “Minimum peak amplitude” was set to “0,” and the “minimum value of maxima” was set to the previously identified threshold. The number of peaks corresponds to the number of axon profiles at that ROI **(Figure 1G-H)**. This was repeated for all ROIs in each nerve section. The number of axon profiles at each ROI was then divided by the number of axon profiles 250 µm proximal to the injury site to calculate the regenerative index. If an image had too much background noise or uneven background brightness, then axon profiles were counted manually. Two to three nerve sections per animal were analyzed. Technical replicates of regenerative indices were averaged for each biological replicate.

### Tyrosine hydroxylase at the neuromuscular junction

The presence of TH, the rate-limiting enzyme of norepinephrine synthesis,^46^ at NMJs reinnervation was assessed in the gastrocnemius muscles at 10 weeks post-injury **(Figure 1E)**.^46^ Gastrocnemii were harvested from perfused mice and cryoprotected with 20% sucrose with sodium azide and allowed to sink at 4°C overnight. Afterwards, the muscle was cryosectioned at 30 µm, and sections were placed on charged glass slides. Gastrocnemii from the contralateral leg of 7 randomly selected mice, which did not receive any intervention or injury, were used as an additional control.

The slides were permeabilized with 0.1% Triton®-X-100 (Fisher BioReagents, BP151-500) in 1X PBS then blocked for 2 hours at RT with 2% bovine serum albumin (BSA, Sigma-Aldrich, A2153), 0.1% Triton in 1X PBS. Rabbit anti-TH (1:500) was diluted in the blocking buffer, and the primary antibody was allowed to incubate for 2 nights at 4°C in a humidity chamber. After 3 10-minute washes with 1X PBS, secondary antibody goat anti-rabbit Alexa Fluor™ 488 (1:200, Invitrogen, A11008) was diluted in the blocking buffer and allowed to incubate at RT for 2 hours followed by 3 10-minute washes with 1X PBS. Following the washes, α-bungarotoxin (α-BTX) Alexa Fluor™ 555 (1:500, Invitrogen, B35451) was diluted in 1X PBS and placed on the slides for 30 minutes followed by 3 additional 10-minute washes with 1X PBS. After the slides were dried, they were mounted with Fluoro-Gel with Tris buffer and imaged on a Nikon Ti-E fluorescent microscope. A 60x oil objective was used with the following exposure times: TRITC 200 ms, FITC 80 ms. NMJs were identified by presence of α-BTX, and the presence of TH was identified by whether or not TH overlapped with α-BTX. At least 50 NMJs were evaluated per mouse with an average of 56.75 NMJs. The number of NMJs with sympathetic innervation was divided by the total number of NMJs evaluated per mouse to calculate a percentage of NMJs that were TH+. The total number of NMJs analyzed was 1,816.

### Retrograde labeling

Nine weeks and 4 days after nerve transection and repair, Cholera Toxin Subunit B, Alexa Fluor™ 555 retrograde fluorescent tracer (CTB555, Invitrogen, C22843) was injected into the reinnervated gastrocnemius muscle **(Figure 1D)**. Each head of the gastrocnemius received 2 µL of tracer delivered via a 26-gauge Hamilton syringe. These would mark the cell bodies of sympathetic neurons whose axons had successfully regenerated to that muscle target. The injections were made in at least two locations in both the medial and lateral heads of the gastrocnemius.

Three days after tracer injection, mice were perfused as described above. Lumbar sympathetic ganglia L2-L5 were dissected from the level of the left renal artery to the bifurcation of the abdominal aorta and mounted onto glass slides with Fluoro-Gel mounting media with Tris buffer **(Figure 1F)**^48^. Images were taken with a 20x objective, 100 ms exposure time on the TRITC channel, with a Nikon Ti-E fluorescent microscope. Labeled sympathetic neurons were identified if the retrograde label filled the soma and extended into the proximal dendrites and if a nuclear shadow could be visualized. Labeled profiles that did not meet these criteria were not counted.

### Statistical analysis

Sample sizes were predetermined based on previous work.^11,45,49,50^ Data are represented as mean ± standard error of the mean (SEM) or as medians with 95% confidence intervals (CI). GraphPad Prism was used to perform statistical analyses (GraphPad Software, San Diego, California USA, www.graphpad.com). Biorender.com was used to create portions of Figure 1.

For the 1- and 2-week acute axon regeneration experiments, 2-way repeated measures analysis of variance (ANOVA) was used to analyze the data followed by Tukey’s multiple comparisons post hoc test. For NMJ reinnervation analysis, one-way completely randomized ANOVA was followed by post hoc Tukey’s multiple comparisons test. Due to the nonparametric nature of cell counts and uneven variances, Kruskal-Wallis test was used to analyze retrogradely-labeled sympathetic neurons. * or + p < 0.05, ++ p < 0.01, *** p < 0.001.

## RESULTS

### Sympathetic axon regeneration into a nerve graft is acutely inhibited by CES

To test the effect of CES on acute sympathetic axon regeneration into a nerve graft, the sciatic nerve received either sham CES, 10-minute CES, or 60-minute CES 1 week prior to nerve transection and repair with a graft **(Figure 1A-C)**. At 1 week post-injury, the regenerative indices were significantly decreased with CES compared to sham CES (2-way RM ANOVA, experimental group factor F(2,12) = 4.219, p = 0.040966, post hoc Tukey’s multiple comparisons test) starting at 1000 µm distal to the injury site (1000 µm sham vs 1-hr p = 0.034888, 1500 µm sham vs 1-hr p = 0.000718, 2000 µm sham vs 1-hr p = 0.028351) **(Figure 2A-D)**. The effect is greater with 60-minute CES compared to 10-minute CES, with 10-minute CES only being significantly different from sham CES at 1500 µm (sham vs 10-min p = 0.009606) and 2500 µm (sham vs 10-min p = 0.036377). Importantly, motor and sensory regeneration was also not enhanced with CES (2-way RM ANOVA, experimental group factor F(2,12) = 1.198, p = 0.3353) **(Figure 2E-H).**

**Figure 2:**
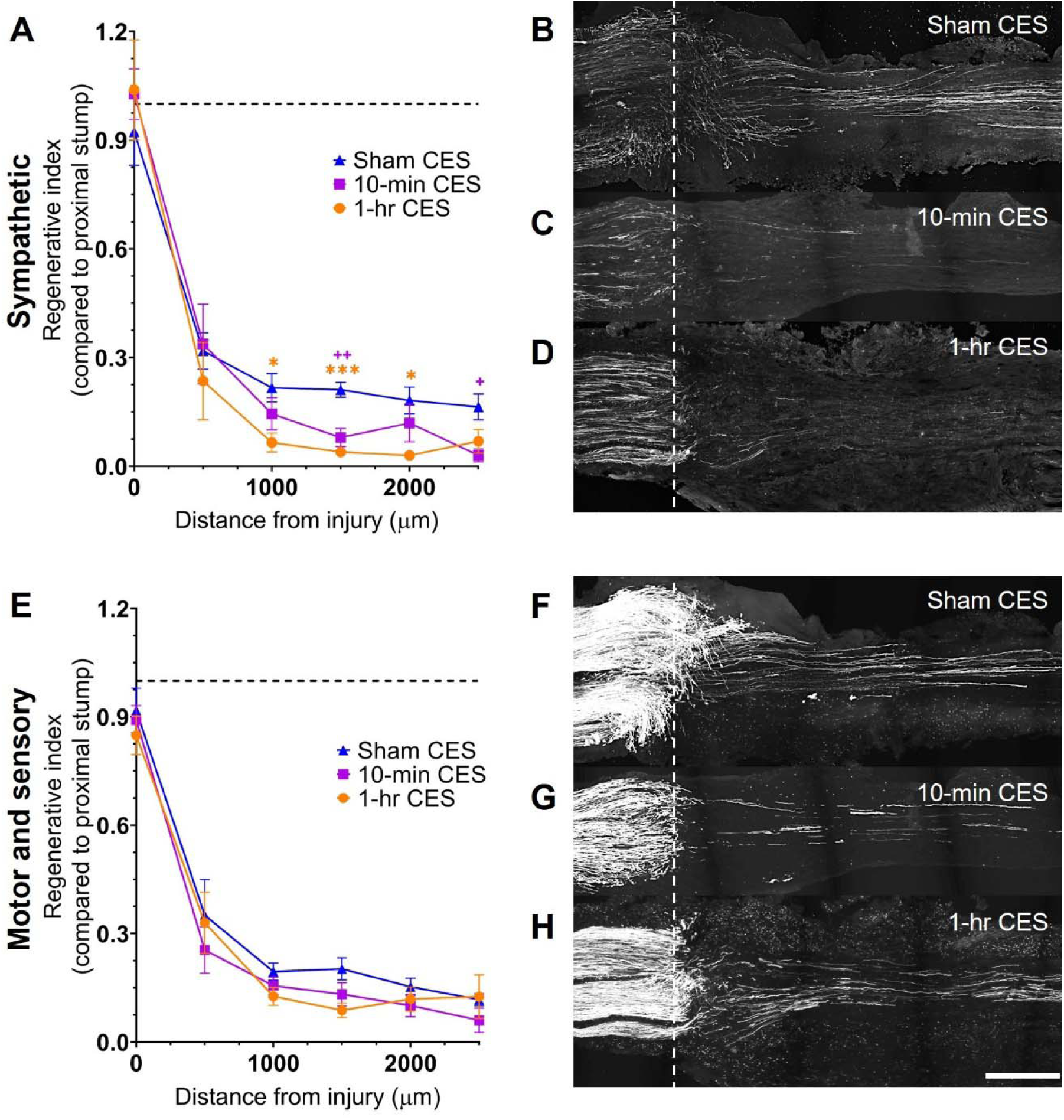
Conditioning electrical stimulation reduces sympathetic axon growth acutely into a nerve graft 1 week after injury. (A) Plot of regenerative indices of sympathetic axons calculated based on number of axon profiles present every 500 µm distal to the injury site relative to the number of axon profiles 250 µm proximal to the injury site. Two-way repeated measures ANOVA with post hoc Tukey’s multiple comparisons test. * = Sham CES vs 1-hr CES, + = Sham CES vs 10-min CES. * or + p < 0.05. ++ p < 0.01. *** p < 0.001. (B) Representative sciatic nerve section after sham CES, (C) 10-min CES, and 1-hr CES with axons visualized with tyrosine hydroxylase immunostaining. (E) Plot of regenerative indices of motor and sensory axons with representative sciatic nerve sections after (F) sham CES, (G) 10-min CES, and (H) 1-hr CES with axons visualized with a yellow fluorescent protein reporter. White dotted line indicates the injury site. Data shown as mean ± SEM. Sham n = 5, 10-min n = 5, 1-hr n = 5. Scale bar = 500 µm.

### Sympathetic axon regeneration after direct nerve repair is not enhanced by CES

Acute axon regeneration in response to CES was also evaluated in the context of a direct nerve repair 2 weeks after injury. At this timepoint, there was no difference in the regenerative indices of sympathetic axons in all groups (2-way RM ANOVA, experimental group factor F(2,17) = 0.7048, p = 0.) **(Figure 3A-D)**. In contrast, motor and sensory axon regeneration was enhanced by 60-minute CES but not by 10-minute CES (2-way RM ANOVA, experimental group factor F(2,17) = 5.465, p = 0.0147) at 1000 µm (1-hr vs sham p = 0.1600, 1-hr vs 10-min p = 0.0418), 2000 µm (1-hr vs sham p = 0.0428, 1-hr vs 10-min p = 0.0307), and 2500 µm distal to the injury site (1-hr vs sham p = 0.2261, 1-hr vs 10-min p = 0.0310) **(Figure 3E-H)**.

**Figure 3:**
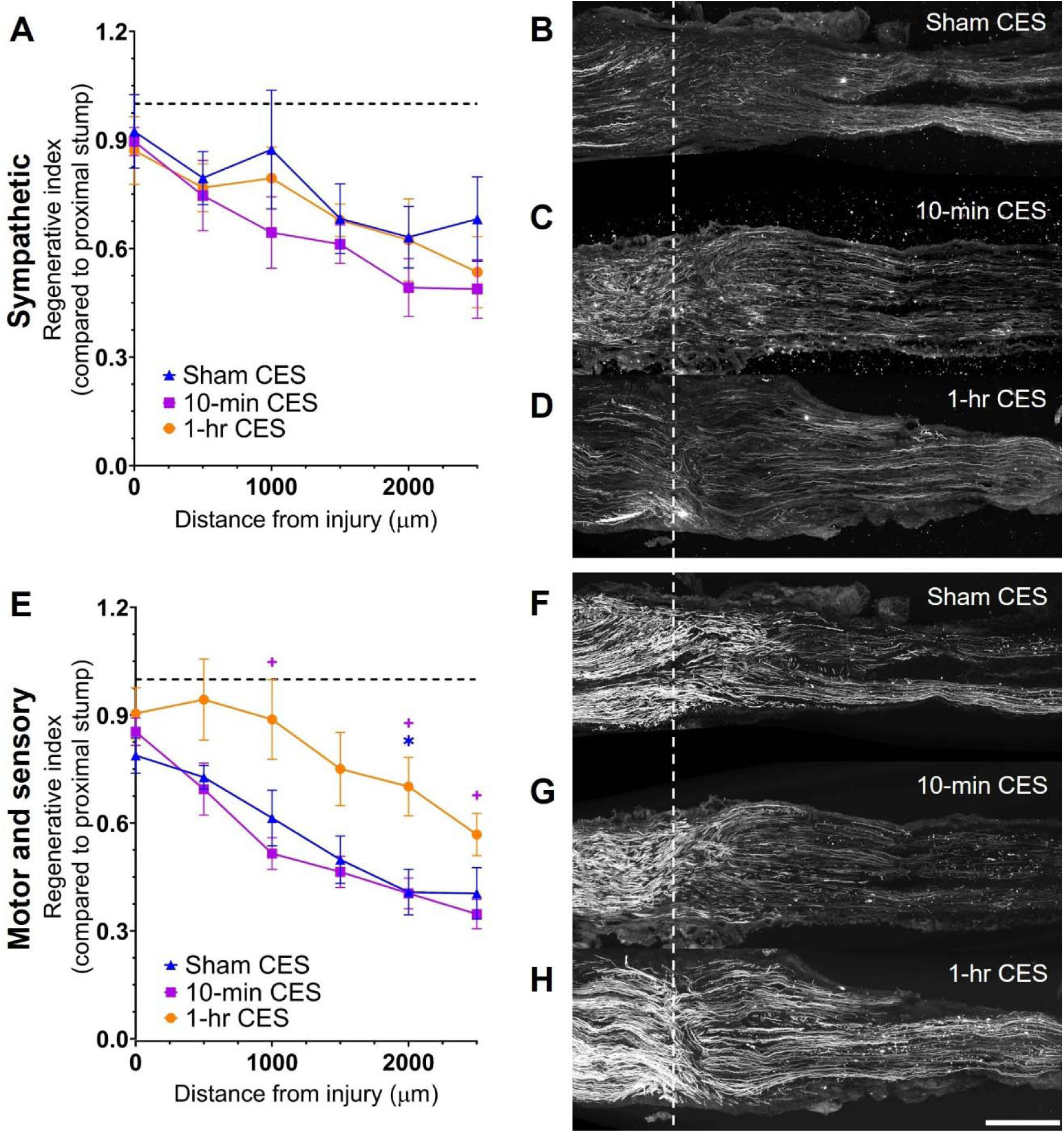
Conditioning electrical stimulation does not enhance sympathetic axon regeneration 2 weeks after direct end-to-end nerve repair but does enhance motor and sensory axon regeneration. (A) Plot of regenerative indices of sympathetic axons calculated based on number of axon profiles present every 500 µm distal to the injury site relative to the number of axon profiles 250 µm proximal to the injury site. (B) Representative sciatic nerve section after sham CES, (C) 10-min CES, and 1-hr CES with axons visualized with tyrosine hydroxylase immunostaining. (E) Plot of regenerative indices of motor and sensory axons with representative sciatic nerve sections after (F) sham CES, (G) 10-min CES, and (H) 1-hr CES with axons visualized with a yellow fluorescent protein reporter. Two-way repeated measures ANOVA with post hoc Tukey’s multiple comparisons test. * = Sham CES vs 1-hr CES, + = 10-min CES vs 1-hr CES. * or + p < 0.05. White dotted line indicates the injury site. Data shown as mean ± SEM. Sham n = 7, 10-min n = 7, 1-hr n = 6. Scale bar = 500 µm.

### Long-term sympathetic reinnervation after injury is unaffected by CES

To evaluate sympathetic reinnervation of peripheral targets 10 weeks after PNI, the sciatic nerve received either sham CES, 10-minute CES, or 60-minute CES prior to transection and repair **(Figure 1A-B)**. A typical NMJ in intact muscle has robust TH expression **(Figure 4B)**, while reinnervated tissue exhibits fragmented NMJs with incomplete TH expression **(Figure 4C)** or negligible TH expression **(Figure 4D)**.

**Figure 4:**
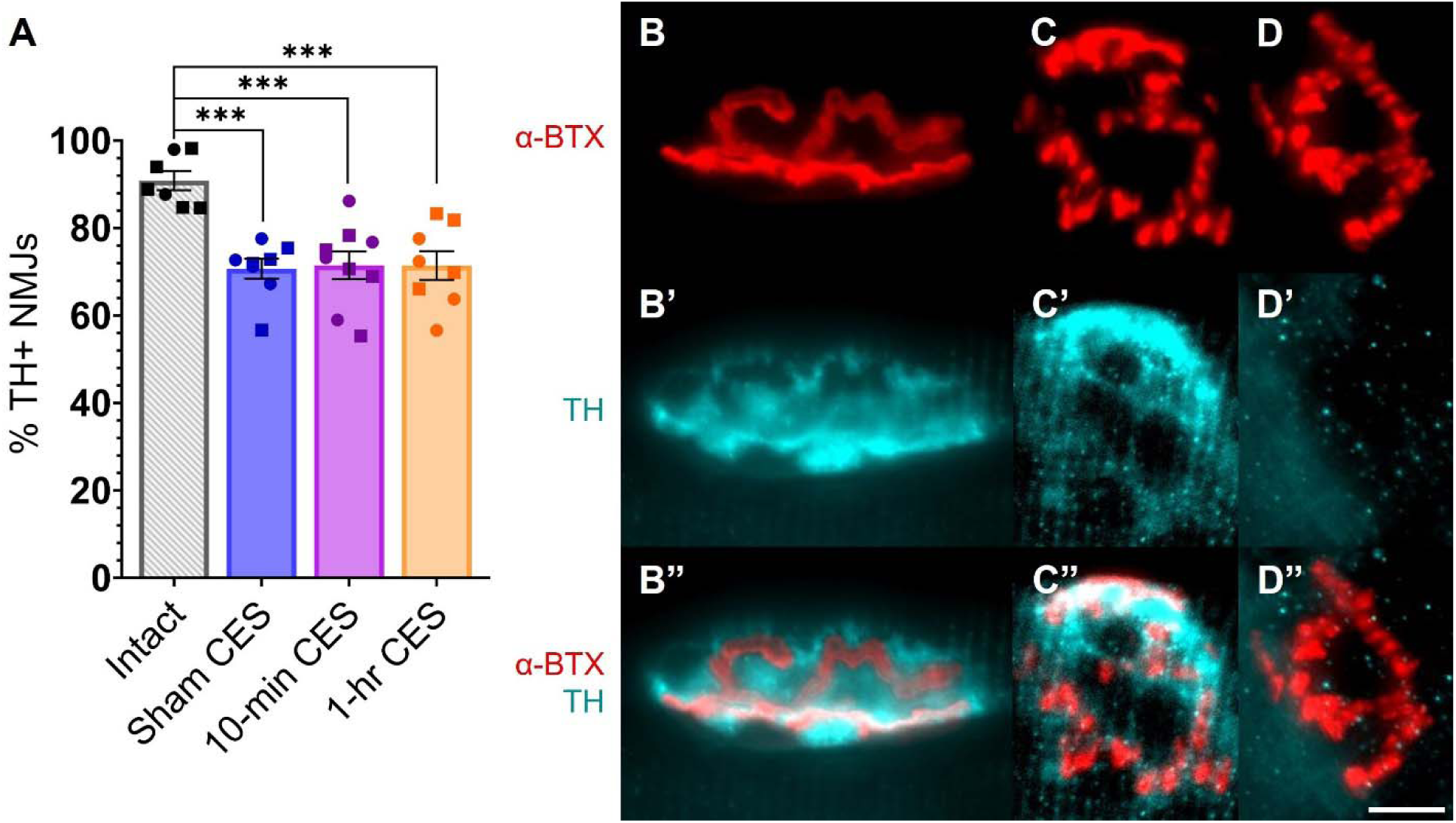
CES does not recover the presence of tyrosine hydroxylase at the neuromuscular junction (NMJ). (A) Percentage of NMJs in the gastrocnemius positive for tyrosine hydroxylase compared to uninjured (Intact) muscles. One-way completely randomized ANOVA with post hoc Tukey’s multiple comparisons test. ***p < 0.001. Data shown as mean ± SEM. Intact n = 7, Sham n = 8, 10-min n = 9, 1-hr n = 8. (B-B”) NMJ from intact muscle with the presence of TH. (C-C”) Fragmented NMJ post-PNI in the gastrocnemius muscle with the presence of TH. (D-D”) Fragmented NMJ post-PNI in the gastrocnemius without the presence of TH. α-BTX = α-bungarotoxin, TH = tyrosine hydroxylase. Scale bar = 10 µm.

The percentage of NMJs that contained TH 10 weeks post-injury was not improved by CES **(Figure 4A)**. All experimental groups exhibited decreased proportions of NMJs that were TH+ compared to intact controls (1-way completely randomized ANOVA, F(3,28) = 10.84, p = 0.000067). The TH+ percentage for all CES groups, including the sham CES group, was 71.2% compared to 90.9% for the intact control group (post hoc Tukey’s multiple comparisons test, intact vs sham p = 0.000244, intact vs 10-min p = 0.000291, intact vs 1-hr p = 0.000388). The proportion of NMJs that was TH+ in the intact control muscles, 90.9% **(Figure 4A)**, is comparable to previous studies.^34^

The L2-L5 lumbar sympathetic ganglia traverse the distance between the aortic bifurcation into the iliac arteries to the level of the left renal artery **(Figure 5A)**.^48,51^ The large L2 lumbar sympathetic ganglia can be seen directly under the diaphragmatic crura **(Figure 5B)**. The ganglia could then be mounted directly to a slide and visualized in its entirety **(Figure 5C)** with clear identification of labeled sympathetic neurons **(Figure 5D)**. The number of retrogradely-labeled sympathetic neurons from the gastrocnemius in the L2-L5 lumbar sympathetic ganglia (shown as medians with 95% confidence intervals) was significantly decreased with 10-minute CES compared to sham CES (Kruskal-Wallis test, p = 0.0218, sham vs 10-min p = 0.0241). No difference was observed between the 2 CES groups (10-min vs 1-hr p > 0.9999) while the difference in retrogradely-labeled sympathetic neurons between the sham CES and 1-hr CES groups was not significant (p = 0.1245) **(Figure 5E)**. The median number of retrogradely-labeled neurons in the sham CES group (287, 95% CI [249.6, 339.2]) exceeded that of both 10-minute CES (194, 95% CI [152.5, 270.6]) and 60-minute CES (222, 95% CI [183.7, 282.8]) groups. The mean rank difference between sham CES and 10-minute CES was 9.479, and the mean rank difference between sham CES and 60-minute CES was 7.500.

**Figure 5:**
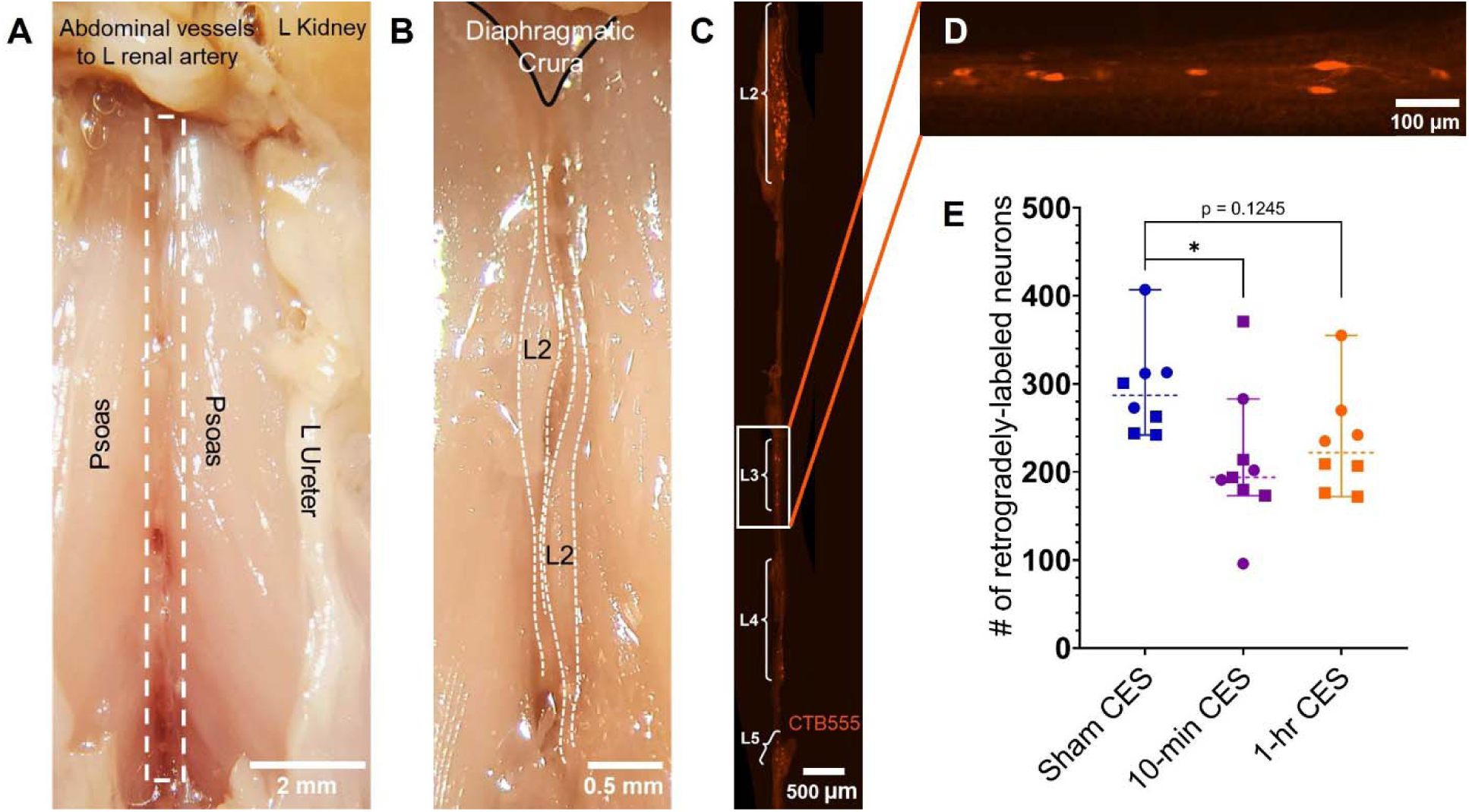
The number of retrogradely-traced sympathetic neurons is not affected by conditioning electrical stimulation. (A) Gross dissection of the lumbar sympathetic ganglia in a perfused mouse. (B) Magnified view of the bilateral L2 lumbar sympathetic ganglia. (C) Unilateral L2-L5 lumbar sympathetic ganglia with neurons retrogradely-labeled with cholera toxin subunit B Alexa Fluor 555 (CTB555). (D) Magnified image of the L3 ganglion with retrogradely-traced neurons. (E) Scatterplot of retrogradely-labeled neurons. Kruskal-Wallis test with Dunn’s multiple comparisons test. * p < 0.05. Data shown as medians with 95% confidence intervals. Sham n = 8, 10-min n = 9, 1-hr n = 8.

## DISCUSSION

CES is a clinically appealing treatment to enhance the regeneration of injured peripheral nerves. In animal models of PNI, CES promotes sensory and motor axon regeneration after several types of nerve repair.^19,21^ In an effort to refine protocols that would further minimize the time needed for nerve stimulation, a 10-minute PES protocol has also been demonstrated to provide similar benefits as a 60-minute PES protocol after nerve injury.^26,27^ Herein, we report the first examination of the effect of CES (10 versus 60 minutes) on the regeneration of motor, sensory, and sympathetic axons.

Sympathetic axon regeneration was acutely inhibited by CES 1 week post-PNI through a nerve graft. The inhibition was more dramatic with a longer application of CES (60 minutes). Intriguingly, CES did not promote motor and sensory axon regeneration through a nerve graft 1 week post-PNI. However, this result could be due to the time point used in this experiment as previous models had evaluated axon growth into a nerve graft at 2 weeks post-PNI.^19,21^ Nonetheless, given that sympathetic outgrowth was impaired with 60 minutes CES, the nerve graft + CES paradigm should be mechanistically explored and independently tested.

With direct nerve repair at 2 weeks post-PNI, no difference was found in sympathetic regeneration between the experimental groups, though it is possible that there were differences in the regenerative front at distances greater than 2500 μm distal to the injury site. At 2 weeks post-PNI with direct repair, 60-minute CES, but not 10-minute CES, was able to enhance motor and sensory axon regeneration, consistent with previous studies.^24^ Therefore, 10-minute CES does not have the same pro-regenerative effects as its 60-minute counterpart.

Since there were no differences in sympathetic regeneration acutely with direct repair, we next evaluated long-term regeneration of sympathetic axons in muscle. The return of functional sympathetic reinnervation of muscle should be a priority because NMJ integrity and muscle mass relies on sympathetic signaling.^34–37^ However, at a longer survival time post-injury with direct nerve repair, there was no difference in the return of TH immunoreactivity at the NMJ between the experimental groups, which is consistent with the acute findings at 2 weeks post-PNI. Furthermore, there were fewer sympathetic neurons that had reinnervated the gastrocnemius with 10-minute CES compared to sham CES. This effect size was comparable to that seen between sham CES and 60-minute CES; however, the difference between 10-minute and 60-minute CES was not statistically significant. Overall, our main conclusion is that CES fails to enhance the regenerative ability of sympathetic axons at both acute and longer times post-injury.

Our study uses the current clinical paradigm of ES that has reached patient populations: 60 minutes of stimulation at 20 Hz with 0.1 ms pulses at relatively low stimulus intensities, which increases the clinical relevance of our experiments. Sympathetic regeneration may not be enhanced by this paradigm because ES recruits the largest diameter, myelinated axons into activity first due to the presence of more parallel ion channels and nodes of Ranvier.^52–56^ The smaller-caliber unmyelinated fibers, such as sympathetic axons, are recruited last and require much higher stimulus intensities and longer pulse durations,^53,57–59^ which may be out of the realm of patient comfort.^60^ ES can also activate surrounding nonneuronal cells such as macrophages and Schwann cells,^61^ which can impact regeneration. For example, ES promotes the release of nerve growth factor from Schwann cells, and sympathetic axons respond to nerve growth factor by sprouting.^32,62,63^ Thus, the current clinical paradigm may not be recruiting sympathetic axons into activity, but sympathetic axons may respond to ES-dependent factors from glia that are released into the regenerative pathway, which could account for the difference in sympathetic axon growth. Although less likely, it is also possible that an initial increase in higher-caliber axonal regeneration outcompetes the smaller-caliber sympathetic axons for available endoneurial tubes, but they eventually find regenerative pathways at later time points. McQuarrie et al. (1978) also observed a “lag” in the regenerative front of sympathetic axons following a conditioning lesion to the sciatic nerve, while others have found that sympathetic axons react to ES with sprouting rather than elongation.^30–32^

In contrast, other studies on superior cervical ganglia have shown enhancement of sympathetic neurite outgrowth in response to a conditioning lesion.^64,65^ Our results appear to be incongruous with those found in the superior cervical ganglia, but congruous with those found in the sciatic nerve,^29^ likely due to differential molecular profiles amongst sympathetic neurons at various anatomical locations.^66^ Therefore, the regenerative ability of neurons in the lumbar sympathetic ganglia, where the postganglionic sympathetic neurons that contribute axons to the sciatic nerve reside, is not comparable to that of the superior cervical ganglia. This underscores the importance of the model chosen and highlights the heterogeneity of sympathetic neurons, which historically were viewed as homogeneous with one common outgoing response.^67,68^ It is increasingly apparent that sympathetic neuron heterogeneity and developmental complexity lead to differences in their behavior, targets, and injury response.

One limitation of our acute regeneration studies is that TH is also present in C-low threshold mechanoreceptors (C-LTMRs), which is a small population of dorsal root ganglia neurons that synthesize and use dopamine or L-DOPA as neurotransmitters.^69,70^ C-LTMRs normally process innocuous mechanical and cooling sensations as well as affective touch.^71,72^ This concern could possibly be circumvented by a primary antibody against dopamine β-hydroxylase (DBH), which converts dopamine to norepinephrine, the main neurotransmitter for postganglionic sympathetic neurons.^73^ The dorsal root ganglia neurons that express TH do not express DBH.^70^ However, our current immunohistochemistry protocols have not allowed for clear visualization of DBH-positive axons in the sciatic nerve, which may be due to the relatively lower amount of *DBH* mRNA present in the distal axons compared to *TH* mRNA.^74,75^ This limitation primarily affects the short-term regeneration assay; therefore, utilizing counts of retrogradely-labeled neurons in the lumbar sympathetic ganglia allowed for a more specific measure of sympathetic regeneration without the confound of neurons from the dorsal root ganglia. The combination of these assays bolsters our main conclusion that CES fails to promote sympathetic axon regeneration, which is consistent with our extensive analysis of motor and sympathetic regeneration after PES.^37^

A strength of this study was the use of a nerve graft for the 1-week acute regeneration experiments, which more closely parallels clinical scenarios in which CES would be used: delayed nerve repair surgery or cases of planned nerve injury.^24^ These delayed surgeries would often involve the use of autologous nerve grafts or nerve transfers,^76,77^ such as a spinal accessory nerve to suprascapular nerve transfer to reanimate the shoulder in brachial plexus palsy,^78^ which is more analogous to the use of a nerve graft in mice. Additionally, CES could be beneficial in situations involving a planned oncological resection.^23,79^ Unfortunately, niether CES paradigm promoted motor, sensory, or sympathetic regeneration through a graft at 1 week post-PNI.

For novel therapeutics to enhance the regeneration of all neuron types more efficiently, including sympathetic neurons, future study directions could focus on altering the ES parameters to better enhance sympathetic growth, a combinatorial ES paradigm to separately enhance motor and sensory growth versus sympathetic growth, or a combinatorial treatment with ES to target genes or proteins that may enhance the growth of sympathetic neurons. Enhancing sympathetic regeneration following PNI is critical for the return of homeostatic mechanisms in the vasculature, skin, muscle, bone, and all other targets of the denervated organs.^34,80–83^ Additionally, aberrant sympathetic outgrowth in the form of sprouting has been implicated in many neuropathic pain states.^31,84,85^ Therefore, exploring approaches that can be combined with ES in a complementary fashion to optimally enhance the regrowth of all neurons will be a critical step in improving patient outcomes after PNI.

## CONCLUSIONS

Despite the success of CES in enhancing the regeneration and functional recovery of motor and sensory axons after PNI,^19,23,24^ both 10 minutes and 60 minutes of CES fail to enhance sympathetic axon elongation and fail to improve sympathetic reinnervation of distal tissues. Importantly, although 10 minutes of PES has been shown to be as effective as 60 minutes of PES at enhancing motor and sensory regeneration,^26,27^ 10 minutes of CES is not as effective as 60 minutes of CES at enhancing motor and sensory axon regeneration. CES is still a valuable therapeutic option that allows for a nerve regeneration intervention to be performed at the bedside prior to delayed nerve repair surgeries and cases of planned nerve resection. However, there remains an urgent need to develop novel therapeutic options that enhance the regeneration of all axon types, which will be an essential step to further improve patient outcomes.

## ACKNOWLEDGEMENTS

This project was supported by the NIH National Institute of Neurological Disorders and Stroke under award number K01NS124912 and in part by a developmental grant from the NIH-funded Emory Specialized Center of Research Excellence in Sex Differences U54AG062334, the Medical Scientist Training Program of Emory University School of Medicine, and the National Institute of General Medical Sciences predoctoral training grant in Genetics T32GM008490. This work was also supported by the Emory University Integrated Cellular Imaging Core Facility (RRID:SCR_023534). The content is solely the responsibility of the authors and does not necessarily reflect the official views of the National Institute of Health. Thank you to Brittney Ward, graduate student, for reading the final manuscript to ensure clarity.

